# Harnessing foresters’ engagement for climate change adaptation: the emerging tool of next-generation citizen science

**DOI:** 10.1101/2025.09.15.676198

**Authors:** Marjorie Bison, Nicole Ponta, Daniella Maria Schweizer, Katalin Csilléry

**Author notes:** **Correspondence** Katalin Csilléry < >. These authors have contributed equally (Marjorie Bison, Nicole Ponta, Daniella Maria Schweizer).

## Abstract

Citizen science is an increasingly common approach for collecting large amounts of data across extensive spatial and temporal scales in ecology and environmental sciences. To facilitate participation, the majority of citizen science projects are based on opportunistic smartphone-based tasks that can be completed in just a few minutes. We reviewed 639 citizen science projects and coordinated distributed experiments, assessing the level of engagement expected from participants, and found that citizens can also perform highly engaging tasks, including conducting experiments, similar to those expected from collaborating scientists. We coined the term “next-generation citizen science” for experiments conducted by citizens with specialized knowledge and addressed the benefits and risks of such projects using the example of MyGardenOfTrees. This unprecedented continent-wide transplant experiment involves over 300 forester-citizens who dedicate their time and expertise to testing different seed origins in their forests over a five-year period, and co-develop a prediction tool with researchers for selecting suitable species and provenances for assisted migration. We used marketing analysis of recruitment data to identify strategies for participant recruitment and retention across the multi-cultural landscape of Europe, thereby expanding the boundaries of citizen science beyond its traditional use. Furthermore, we present the development of the experimental design and protocols aimed at optimizing logistical feasibility, scientific rigor, and social acceptance. Our findings highlight the untapped potential of such experimental citizen science approaches to increase the scale of ecological experiments and ultimately obtain generalizable findings, thereby overcoming context dependence.

## 1 INTRODUCTION

The quest for general patterns and predictable processes in nature has been a perpetual goal of ecologists and evolutionary biologists (Mc Arthur and Steeves 1972, Shrader-Frechette and McCoy 1994). This quest has never been more relevant than it is today, given the ongoing global changes causing unprecedented biodiversity loss. However, context dependency arising from interactions between biological processes and the environment often leads to findings that are not generalizable and therefore not transferable among sites or organisms (Catford et al. 2022). While ecologists worldwide agree that there is an urgent need to increase the spatial and temporal scale of studies, fundamental challenges exist related to funding, project management, and academic barriers, such as the pressure to produce rapid results (Kuebbing et al. 2018). To overcome the limitations of single-site and/or short-term studies, systematic reviews and meta-analyses have been proposed (Arnqvist and Wooster 1995), and more recently, entire new research fields, such as macroecology (Gaston and Blackburn 2000) and macrogenetics (Leigh et al. 2021), have been established to mine data for general patterns.

Although these approaches have been fundamental in advancing the synthesis of case studies into generalizable knowledge (e.g. Shade et al. (2018)), their power is often limited due to differences in design and methods across studies (Fraser et al. 2013, Knapp et al. 2012, Cocciardi et al. 2024).

The scale of experiments can also be increased when collaborating research teams run experiments at multiple locations while following the same protocol, a so-called coordinated distributed experiment (CDE) Fraser et al. (2013), Borer et al. (2014). Indeed, several CDEs (even though they have not consistently been named as such), such as long-term ecological networks (LTERs) or several international provenance trials in forestry (e.g. (Liepe et al. 2024)), have led to considerable advances in our understanding of processes behind patterns (Borer et al. 2014, Knapp et al. 2012, Cocciardi et al. 2024), such as litter decomposition (Woodward et al. 2012, Gholz et al. 2000), nutrient dynamics (collaborators 2014, Keller and Taylor 2008), local adaptation and evolutionary potential of forest trees (Alberto et al. 2013). Citizen science (CS), broadly defined as the partnering of scientists with volunteers to answer scientific questions (Dickinson et al. 2012), is also increasingly used to reach larger spatial and temporal scales of observational studies (Bonney et al. 2009), particularly in monitoring the environment and biodiversity research (Theobald et al. 2015, Johnston et al. 2023, Kobori et al. 2016, Soroye et al. 2018, Pocock et al. 2018). In this paper, we investigate the benefits of involving citizens with specialized knowledge in hypothesis-driven research and experimental research, using the forestry sector as an example.

Forest trees are pillars of terrestrial ecosystems, yet are increasingly experiencing die-back related to rapid climate change, deforestation, and urbanization (Shaw and Etterson 2012, Trumbore et al. 2015, Wessely et al. 2024, Senf and Seidl 2021, Allen et al. 2015, Hoffmann and Sgrò 2011, Aitken and Bemmels 2016). Assisted migration (AM), defined here as any climate-change-induced human-aided translocation of species within or outside of their range, is increasingly recognized as a necessary climate change adaptation and conservation strategy Hällfors and Dalrymple (2022). It has been included in several national programs, such as the New EU Forest Strategy 2030 (COM(2021)0572) in Europe. However, several challenges related to context dependency hinder the development of effective AM programs. First, local adaptation may not be universal, particularly at the range margins, where inbreeding, gene swamping, or expansion load may lead to maladaptation (Angert et al. 2020, Valladares et al. 2014, Fréjaville et al. 2020). Second, existing studies often suffer from limited statistical power, due to inadequate representation of populations across species ranges (Blanquart et al. 2013, Gibson et al. 2016). Third, experimental environments, i.e., common garden trials, are frequently manipulated in ways that do not accurately reflect natural conditions in terms of soil, microclimate, and biotic interactions (Gibson et al. 2016, Alexander et al. 2016), limiting our understanding of the relative roles of genetic, environmental factors, and phenotypic plasticity (Schwinning et al. 2022). Ultimately, our understanding of adaptation is primarily based on production traits measured in young adult trees, while neglecting early life-history traits that are crucial for establishing future generations of locally adapted trees (Donohue 2014, Talluto et al. 2017, Isaac-Renton et al. 2018). Considering the genetic correlations among traits is essential for anticipating species responses to environmental changes (Juenger 2013, Csilléry et al. 2020ab, Climent et al. 2024), particularly in light of novel selection pressures that may affect production traits indirectly (Montwé et al. 2018, Zohner et al. 2016, Stinziano and Way 2017).

The objective of this study is threefold. First, we reviewed over 650 CS and CDE projects and evaluated their evolution in terms of spatial extent and level of engagement required from participants. This analysis enabled us to situate our CS project: a pan-European, thus continent-wide, experimental citizen science project, MyGardenOfTrees (www.mygardenoftrees.eu). This project involves forester-citizens and achieves an engagement level comparable to that assumed in CDEs. MyGardenOfTrees aims to estimate so-called reaction norms of genetic units (Stearns 1989) using a representative set of experiments that explore the multi-dimensional gene–environment space (Fig. 1). In turn, such reaction norms can be used to develop a prediction model (Jarquín et al. 2014) to advise AM decisions, which in turn serves the participating foresters. Our second objective is to showcase the opportunities and barriers to recruiting forester-citizens across the multicultural and linguistic landscape of Europe in MyGardenOfTrees, as well as the experimental design of the project to optimize logistical feasibility, scientific rigor, and social acceptance. Third, we conclude by identifying other research questions in ecology and evolution, where a similar experimental CS would be beneficial, especially in the context of global change.

**FIGURE 1.**
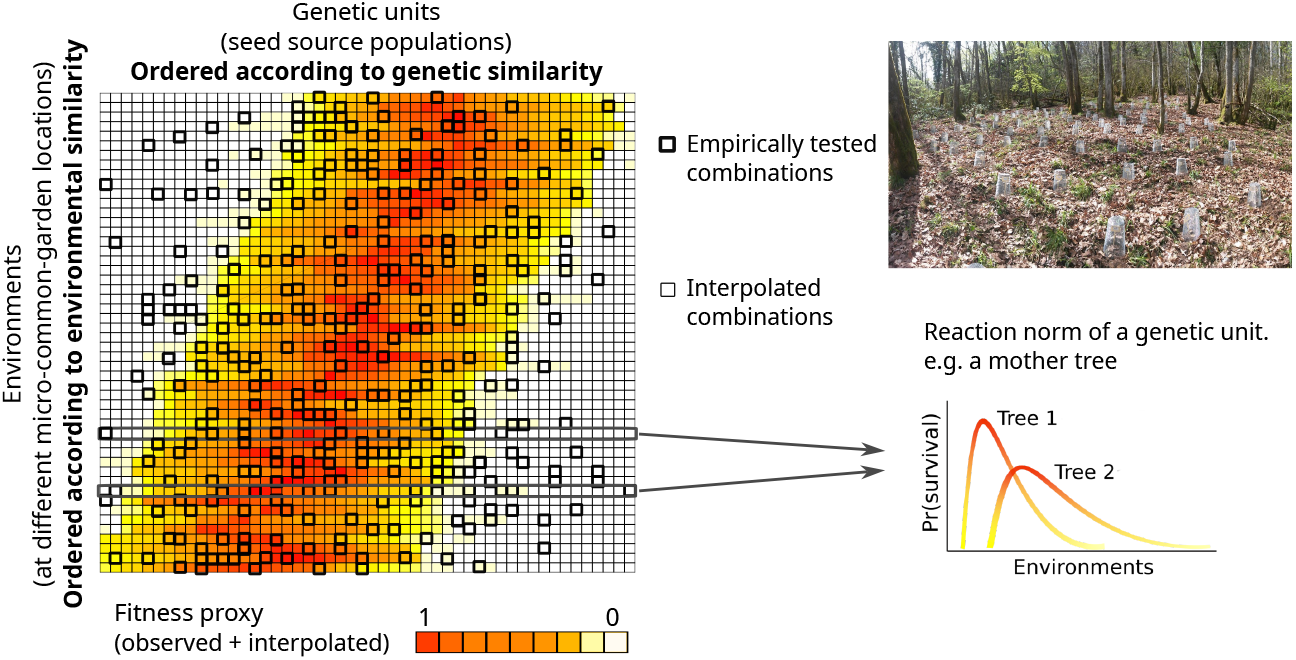
The concept of distributed common garden experiments to obtain generalizable results. (a) Reaction norms of different genetic units vary across different environments. The panel illustrates an ideal case where gene–environment interactions are predictable. (b) Example of an experimental unit, a so-called micro-garden. Seeds are protected from seed predation and the forest site is protected from contamination by foreign seeds using custom-designed and custom-produced seed protectors.

## 2 MATERIALS AND METHODS

### 2.1 Dataset of CS projects and CDEs

We complied a dataset of CS projects from Global (https://scistarter.org), North American (https://www.citizenscience.gov, https://science.gc.ca/site/science/en/citizen-science-portal), Australian (https://biocollect.ala.org.au), Indian (https://citsci-india.org/projects), and European (https://eu-citizen.science, https://data.jrc.ec.europa.eu/dataset, https://www.mitforschen.org, https://www.citizen-science.at, https://www.citizenscience.uzh.ch) websites. We only included projects initiated by and/or involving universities or research institutes, using the “institution type” filter of “academia” in the European databases and funding sources such as “Smithsonian” or “National Science Foundation” in the SciStarter and US Government databases. For the other databases, we searched all environmental projects to select those involving a research institution. We focused on CS projects relevant to ecology and evolution, using only the following keywords: “Ecology and Environment,” “Plants,” “Animals,” “Agriculture,” “Biology,” “Birds,” “Ocean, Water, Marine Terrestrial,” and “Insects and Pollinators.” We entered projects multiple times if they proposed more than one action in which citizens could engage. Our dataset includes 623 CS projects and 16 CDEs (Dataset S1).

We extracted the country where the project was initiated, the start date of the project, and the target public, either from the CS databases searched or directly from the project websites. The term “general public” was used when no selective criteria were given. “Public subset” was used when the project targeted only a specific age class, sex, physical address, or language group. “Public with affinity” was used for projects that included participants with an affinity/sensitivity towards the research question and/or the study species and/or participants who have a hobby that brings them to the correct location and/or time when the observations are being made. “Public with special knowledge” was used when participants had to possess prior knowledge acquired through formal (e.g. farmers, foresters, rangers, or scientists) or informal education (e.g. bird watchers or divers). The project protocol relied on the fact that participants had this previous knowledge. We assessed the level of participant engagement according to a set of simple criteria (Table 1) that could be reliably extracted from the project website or protocol. We calculated an engagement score by assigning a value of one to each fulfilled criterion and summing them up. The lowest score was zero, and the highest was eight. We analyzed the evolution of CS using a linear filter on the time series of the engagement score, target population, and geographic extent from 1978 to 2024, employing a moving average implemented in the “convolution” method in the `stats` package in R (version 4.4.2).

**TABLE 1.** Definition of the engagement score of a citizen science (CS) project or a coordinated distributed experiment (CDE). Each line describes an aspect of engagement. Projects were assigned an engagement score of zero if they did not include these aspects. Each aspect increased the engagement score by one.

| Engagement type | Example |
| --- | --- |
| None | Citizens submit photographs of sharks taken anywhere and on any date<sup>1</sup> |
| Data collected in a specific place | Citizens use online forms to send observations of the environment around Lake Tahoe, USA<sup>2</sup> |
| Data collected in a specific time of the year/week/day | Citizens send night sky brightness observations<sup>3</sup> |
| Data collected in the same place at least twice | Citizens place cookies in their garden and then collect ants on them<sup>4</sup> |
| Commitment for more than one year/season | Citizens are assigned a specific place to monitor a rare species<sup>5</sup> |
| Presence of a written protocol | Citizens follow a protocol for monitoring dragonflies<sup>6</sup> |
| Collection of one data point takes at least 30 minutes | Citizens collect samples to assess water quality of a lake in Missouri<sup>7</sup> |
| Compulsory in-person training | Following in-person training, citizens monitor aquatic insects to assess water quality of a stream<sup>8</sup> |
| Manipulation of specialized equipment | With provided material, citizens build a multi-sensor system to monitor insects<sup>9</sup> |
| Samples are sent/brought to the researchers | Citizens capture and raise caterpillars of monarch butterflies and send a sample to the researchers<sup>10</sup> |
| Manipulation of the environment | Citizens implement a fenced open-field experiment and apply a fertilizer treatment<sup>11</sup> |

<sup>1</sup> https://sharksofcalifornia.fieldscope.org/; <sup>2</sup> https://citizensciencetahoe.org/home; <sup>3</sup> https://globeatnight.org/; <sup>4</sup> https://scistarter.org/ant-picnic; <sup>5</sup> https://plantsofconcern.org/; <sup>6</sup> https://illinoisodes.org/; <sup>7</sup> https://www.lmvp.org/; <sup>8</sup> https://therouge.org/bug-hunts/; <sup>9</sup> https://kinsecta.org/citizen-science/; <sup>10</sup> https://monarchparasites.org; <sup>11</sup> https://nutnet.org/home

### 2.2 Participant recruitment data

Initially, MyGardenOfTrees targeted all European countries for recruitment. However, some countries had to be abandoned early on because the recruitment of a local coordinator was not successful and/or the target species were of limited interest for local forestry (i.e. Baltic and Scandinavian countries, United Kingdom, Greece), or because the country was small (e.g. Andorra, Liechtenstein) or had low forest cover (e.g. Portugal), which was sometimes coupled with a unique official language (e.g. Montenegro). In the target countries that could be assigned to a local coordinator (Table S1), we undertook various advertising activities, subsequently called recruitment actions (Table 2). Every local coordinator reported the recruitment actions they performed across Europe, specifying the target country, language, item type, media type, publication date, and number of people reached. A total of 273 recruitment actions were recorded (complete list in Dataset S2) between July 1, 2022 and March 31, 2024. We measured recruitment as the number of foresters who completed the registration form implemented in LimeSurvey and made available on the project website (see all questions and their order in Text S1). We collected basic socio-economic data, the type of forest for the future micro-garden, and a marketing attribution question asking where the participant had heard about the project. Attrition rates were low; out of 381 foresters who began filling out the registration form, 302 (79%) completed the entire registration process, received the materials, established a micro-garden in the forest, and submitted the first field observation form about the experiment (Dataset S3).

**TABLE 2.** The four main types of recruitment actions used for MyGardenOfTrees by the local coordinators across the 20 participating European countries. Impact indicates the median (maximum) number of people reached by a single action of the type. For outreach, the maximum indicated the total number of visits to the project’s website during the recruitment period. For social media, impact indicates the number of visits to the project’s Facebook page in a single day.

| Action type | Examples | Impact |
| --- | --- | --- |
| Outreach | Website development (by country)<br>Article in magazine for professionals<br>Newspaper article by a journalist<br>Project videos | 100 (2781) |
| Direct contact | General email to a large target group<br>Personalized email to a person or a small target group<br>Phone call to a person<br>Online or personal meeting with a person or small group | 5.5 (250) |
| Events | Unplanned informal chat at events (conferences, fairs)<br>Formal participation at events (talk, stand, poster) | 25 (100) |
| Social media | FaceBook posts<br>LinkedIn posts | 5 (88) |

We used marketing mix modeling (MMM) to estimate the impact of recruitment actions on registrations (see Materials and Methods). Using marketing language, we had a single “product”: participation in MyGardenOfTrees. This was our “sales”, thus the response variable. Our main objective was to determine the most effective recruitment action type, thereby providing information on how to optimize recruitment in a similar project in the future. We described the effect of each action by “marketing reach”, i.e. the number of people who were reached by our actions. We used various proxies to measure this. For outreach recruitment, we utilized Google and YouTube analytics for the project websites and videos, while for articles in newspapers and magazines, we relied on the number of subscribers. For direct contacts, we counted the number of persons directly targeted, such as unique email addresses or participants in video calls, or the number of recipients when the email was sent to a mailing list. For events, we counted the number of outreach materials distributed from the project’s stand, including flyers and printouts. Finally, for social media, we used the number of likes on Facebook and LinkedIn for posts related to the project, as well as the number of followers on the project’s Facebook page. We aggregated both the response variable and the number of people reached by actions to weekly sums, as is usual in marketing analysis.

We used adstock theory (Broadbent 1979) to model how awareness of a product declines over time. For direct contact, events, and social media action types (Table 2), we used a simple decay model:

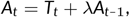

where *A*_*t*_ is the advertising stock at time *t, T*_*t*_ is the advertising spend at time *t*, and lambda is the adstock decay rate. We implemented the function using the `filter` function’s recursive method (R package `stats`). For the recruitment action type, outreach, we attempted to use a delayed decay model (eq. 3 in Jin et al. (2017)), including an adstock decay parameter, *λ*, the delay of the peak effect, *θ*, and the maximum duration of a carryover effect. While the model could be fitted and the parameters estimated (*θ* = 4 weeks, *L* = 6 weeks), a model with a simple decay function for outreach provided a better fit. We jointly estimated the *λ* parameters and the effect sizes of the different recruitment action types using the `nlsLM` function in the R package `minpack.lm`. Analysis scripts are available at https://github.com/kcsillery/MyGardenOfTrees.

Additionally, we conducted a semi-structured interview with 12 local coordinators to gather information on their recruitment strategies and the barriers they encountered (Text S3). The interviews were conducted between March and May 2024, in English, and lasted around 30 minutes. We analyzed transcripts of the interviews using a mixedmethods approach that combines deductive and inductive thematic analysis (Neuendorf 2018), employing MAXQDA (VERBI 2021). The first step in coding the interviews was to use word clouds and word frequency analysis to find emergent themes (themes that were repeated within and across the interviews). We identified three main themes: strategy, barrier, and lessons learned. In the following section, we focus on barriers because they provide complementary information to the analysis of recruitment actions that centered on “what worked.” We classified the barriers into several sub-themes (Dataset S4) and compared them between Eastern, Western, and Southern Europe (Fig. 4d).

### 2.3 Climate and composition of European forests

We used EU-Forest (Mauri et al. 2017) to extract the areas of Europe where (*Abies*) and (*Fagus*) are currently growing, based on harmonized data from national forest inventories. We used these occurrences to establish the climate envelope of the current distribution. From EU-Forest, we also extracted the number of forest tree genera per unit area. We compared it with the number of tree species/genera that the participants reported in their forest, where they planned to establish their micro-garden, using a Spearman rank correlation test. We retrieved data on the proportions of forests that are privately vs publicly owned from FAO’s Global Forest Resources Assessment 2010 report Food and of the United Nations. Forestry Department (Rome).

### 2.4 Experimental design

Seeds were collected from 19 fir (*Abies*) and 14 beech (*Fagus*) provenances, with an average of 10 families per provenance, resulting in a total of 285 genetic units to be tested across 302 environments, represented by the participant micro-gardens. This required the preparation of 302 unique seed parcels, each containing four replicates (to be established as blocks) of 25 unique genetic units (Fig. 1b; Text S2). To minimize errors during parcel preparation and ensure logistical feasibility, we divided the provenances into four groups, each containing five fir and three to five beech provenances. Each block contained three replicates of a provenance represented by different families. We defined the groups to ensure that the main genetic and geographic clusters of the species were described based on our existing knowledge (Liepelt et al. 2009, Magri 2008). Further, we ensured that each participant received an “exotic” species of fir or beech, as there was high interest among foresters in obtaining such seeds. We characterized the climate of the test sites using longitude, latitude, elevation, and the CHELSA dataset and its bioclimatic variables for the time range 1981–2010 (Karger et al. 2017), extracted for each approximate micro-garden location as communicated by the participant on the registration form.

We employed the method of anticlustering to partition a set of units (Feo and Khellaf 1990, Papenberg and Klau 2021, Brusco et al. 2020), specifically different genetic units, into groups represented by the various environments of the micro-gardens. The anticlustering approach is the reverse of clustering analysis, intended to achieve high within-group diversity. We performed anticlustering using the R package `anticlust` on the climatic distance matrix, calculated using the R package `ecodist` on scaled climate variables. We used the diversity criterion option, which maximizes between-group similarity by maximizing within-group heterogeneity, with K=4 representing the number of micro-garden types, and the method of (Brusco et al. 2020) with 5000 repetitions. We then randomly assigned families to the garden types, ensuring each block had 25 unique genetic units and respecting the available seeds. This was done to ensure that participant observations corresponded to a unique seeding spot, while allowing flexibility in the sowing order for establishment of the experiment (Text S2). Note that although the experimental design was conducted using the CHELSA data, we present the environmental space here with the Copernicus ERA5 dataset, as it also includes the year of the experiment, 2024 (Hersbach et al. 2023) (accessed May 1, 2025).

## 3 RESULTS

### 3.1 The emerging tool of next-generation citizen science (NGCS)

Analysis of the 639 CS projects reveals that CS emerged independently in different places and times, with some flagship projects that are still running today (Fig. 2a). For example, the ladybird beetle observation network in Belgium started as early as 1800 (available from the Global Biodiversity Information Facility (GBIF); (GBIF 2024)), the phenology observation network in Austria (Vogelwarte) has been running since 1851 (though interrupted several times), and the Audubon Christmas Bird Count in the USA started in 1900 as a contest to discourage bird hunting. Intimately linked with the development of conservation biology, CS began to gain popularity in the 1980s, especially in economically developed countries in North America and Europe (Fig. 2b). In these early projects, participants were highly engaged based on our list of possible engagement types (Table 1): they were required to follow complex and demanding protocols, repeat observations at specific locations or at particular times, or manipulate specialized equipment (see Dataset S1 for more examples). In contrast, the geographic scale of projects was relatively limited, reflecting the slow communication tools available, such as the post.

**FIGURE 2.**
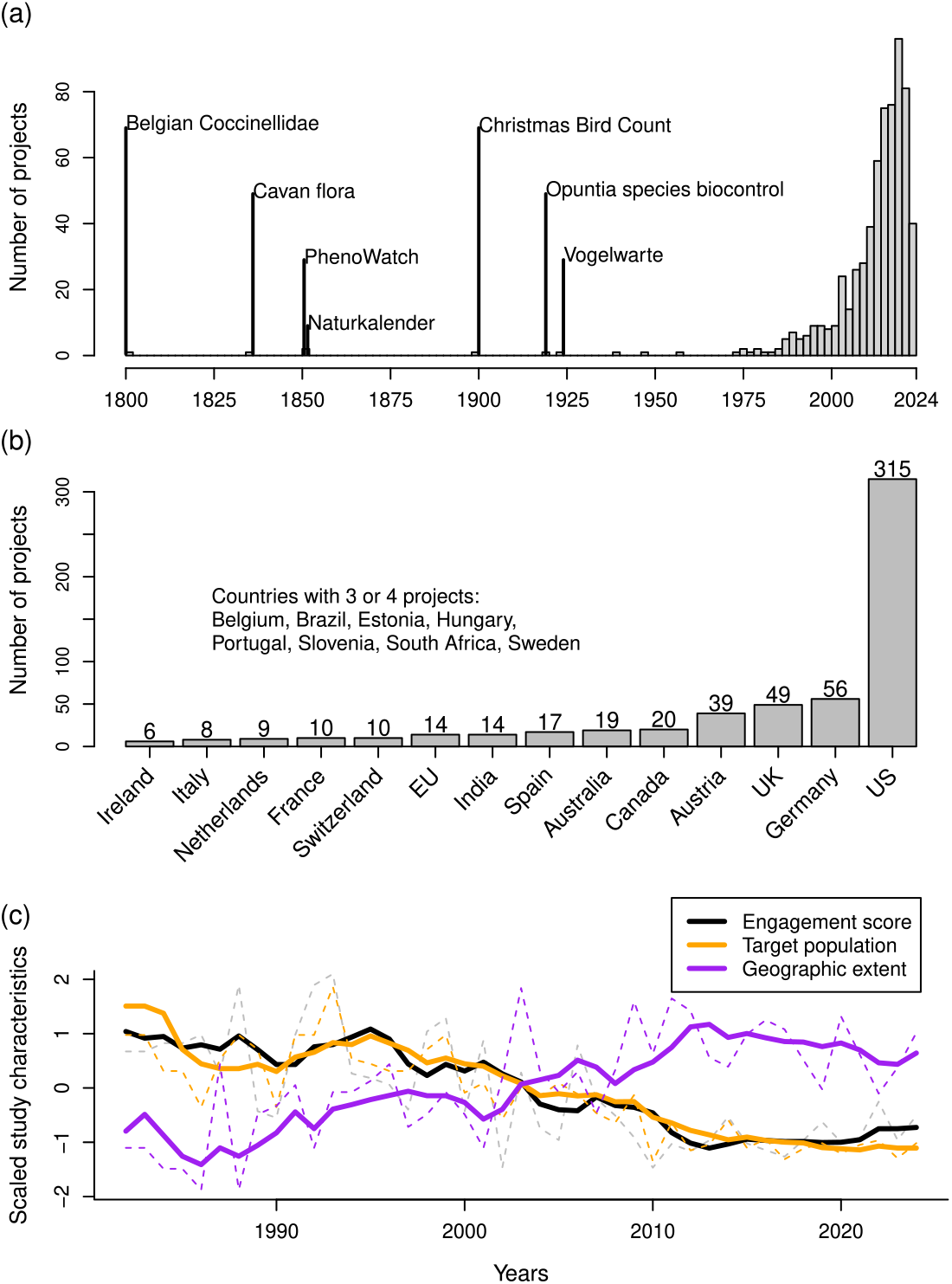
The history of citizen science (CS) in biodiversity research, explored through a novel dataset compiled in this study (see Materials and Methods). (a) The timeline of CS projects, (b) the number of projects per country across the entire considered period, and (c) the trade-off between the geographic extent (ordinal scale: local, subnational, national, regional, continental, global), target population (ordinal scale: general public, public subset, public with affinity, public with specialized knowledge, fellow scientist), and the engagement score (see Table 1 for the definition) of CS projects through time, based on linear filtering with a moving average (see Materials and Methods).

As is often the case, the real breakthrough in CS happened at the moment of significant technological advances: the World Wide Web (2000) and, later, smartphones (early versions from the mid-2000s) revolutionized communication, and enabled CS projects to expand across countries and continents and to reach out to more citizens thanks to the eased data collection and sharing (Fig. 2a and c). In parallel, the creation of simpler numeric protocols, such as taking a photo of any individual plant or animal (e.g., in iNaturalist https://www.inaturalist.org/) or annotating an image from a camera trap (e.g., Snapshot Serengeti https://www.zooniverse.org/projects/zooniverse/snapshot-serengeti), has led to a decrease in the level of engagement required from participants (Fig. 2c). Indeed, in over 30% of projects in our dataset, none of the engagement types listed in Table 1 are relevant. Today, such mass-participation projects, targeting the general public across large spatial scales and requesting opportunistic data collection, are dominating the CS landscape (Fig. 2c). While they have well-recognized benefits, they also have shortcomings related to data quality (Hoeffner et al. 2025, Nov et al. 2014, Anhalt-Depies et al. 2019, Barbato et al. 2021). Indeed, oftentimes, data from such projects are not utilized in scientific publications (Theobald et al. 2015).

Review of CS also highlights that some projects have resisted the current trend, recruiting a specific subset of the population to engage them in complex tasks. In fact, when comparing the engagement scores of CDEs and CSs, we notice that CS projects that work with citizens with specialized knowledge can have participant tasks that are just as engaging as those conducted by scientists participating in CDEs (Fig. 3). The most engaging CS projects involve bird ringing (e.g. https://odnature.naturalsciences.be/bebirds/), butterfly monitoring along transects available for volunteers with appropriate taxonomic skills (e.g. https://www.dagfjarilar.lu.se/), or bat monitoring, where volunteers are trained to use handheld ultrasonic detectors (e.g. http://wiatri.net/inventory/bats/). While most experimental projects rely on the participation of fellow scientists, a handful of CS projects have demonstrated that citizens can even conduct experiments when the tasks align with their profession or special skills. For example, in SurfingForScience.org, citizen surfers manipulate microplastic trawl nets and take water samples. One of the most prominent examples, however, is participatory or collaborative plant breeding, where farmer-citizens test (van Etten et al. 2019) or even breed (Goldringer et al. 2020, Santamarina et al. 2025) agricultural varieties. Such experimental projects require targeted recruitment of a specific subset of the population, potentially those with special skills or knowledge, to conduct high-effort sampling, following a particular protocol (Kobori et al. 2016, Schröter et al. 2017, Castagneyrol et al. 2023, Brown et al. 2019). We coined the term “next-generation citizen science” (NGCS) to refer to experimental approaches that involve collaboration with non-scientist professionals, using state-of-the-art communication and data handling technologies, to address scientific questions highly relevant to the participants’ profession, skill set, or hobby.

**FIGURE 3.**
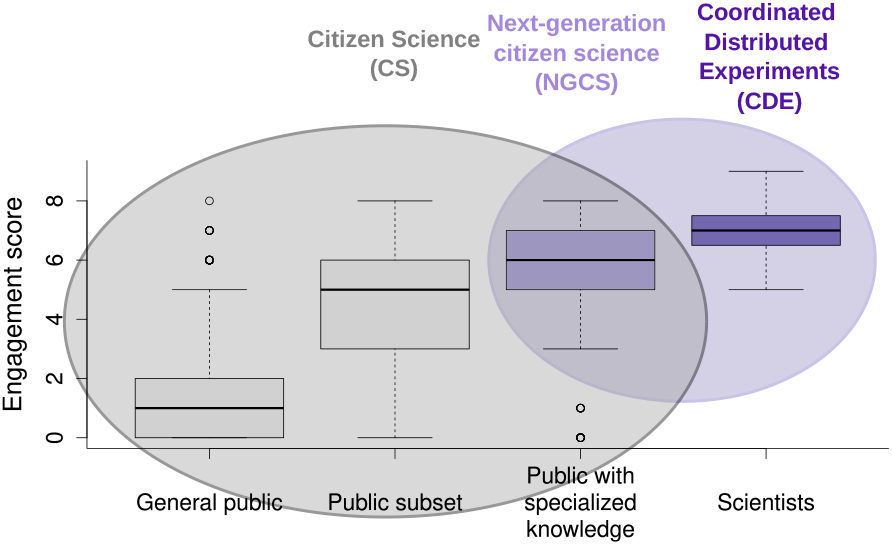
Next generation citizen science (NGCS) at the intersection between citizen science (CS) and coordinated distributed experiments (CDEs). Engagement scores of CS projects in our dataset (see Materials and Methods), categorized by the target population and simplified into three categories. Public with specialized knowledge includes farmers and foresters, for example.

MyGardenOfTrees is an NGCS project that recruited professionals, i.e., European foresters, to carry out an experiment that builds on their professional knowledge and experience (Fig. 2b). The project’s engagement score is eight, and participants are required to follow the experiment over a period of five years. This involves dedicating a 100 m^2^ forest area that they own or manage to the experiment and committing, on average, 35 hours of voluntary work in the first year, with a smaller amount of time in the following years as seedling mortality occurs (Text S1 and S2). The project focuses on two major European tree species, European beech (*Fagus sylvatica* L.) and silver fir (*Abies alba* Mill.), and their Mediterranean and Oriental sister species (*Fagus orientalis* Lipsky, *Abies nordmanniana* (Steven) Spach and *Abies borisii-regis*). Both species complexes have a high economic and ecological importance in Europe, and some are discussed as species resilient to climate change or as potential species for AM (Mauri et al. 2016, Kurz et al. 2023). In the following section, we showcase the lessons learned from MyGardenOfTrees’ recruitment strategy, and we describe how this continent-scale experiment was designed to minimize the pitfalls specific to CS (Prendergast et al. 2022).

### 3.2 Recruitment of foresters across Europe

Participants in CS are not customers, yet there are parallels with customer-facing sectors, such as the need to market a product/project, enhance the customer/participant experience, and retain customers/-participants (Hart et al. 2022). Initially, CS projects can benefit from recruiting strong participants, referred to as “champions” (Jacobs et al. 2005), who will serve as amplifiers of the project’s goals. We employed this strategy during two sets of pilot trials, “2021 summer trials” and “2021–2023 trials” (see https://www.mygardenoftrees.eu/), where we recruited a handful of colleagues, foresters, environmental activists known to us through our network, and even family members. The implementation of these trials and the preliminary results obtained from them were crucial in launching the more ambitious main trials, which aimed to recruit hundreds of forest owners and managers from across Europe. Notably, at that point, we could create the first outreach media items, such as the project website, magazine articles, videos, social media posts, and elaborate on the experimental protocol. In the main “2023–2028 trials”, champions, hereafter referred to as “local coordinators”, were actually hired for the project. They were responsible for recruitment in one or several countries, while also participating, which helped us overcome the omnipresent linguistic and cultural barriers in Europe (Table S1).

We utilized four main recruitment action types for the recruitment of voluntary foresters (Table 2; Dataset S2). First, creating outreach materials was the initial essential step in the entire recruitment process, as we used and referred to these materials in other types of actions. The primary outreach materials were the project website, with 18 country-specific versions, outreach articles, and a project flyer. Second, directly contacting potential participants relied on prior knowledge of contacts, which were obtained from the countries’ national forest service and private owner associations, and were extended through the networking activities of the local coordinators. Third, we attended the principal forestry fair in several countries, where we rented a stand. Finally, we utilized social media to amplify the impact of our outreach actions, such as posting news items on the website. During the recruitment period, we conducted a total of 273 recruitment actions, reaching approximately 16,000 people across 23 countries (Dataset S2).

We used marketing mix modeling (MMM) to estimate the impact of each recruitment action type on the registrations, using the number of people reached by each action as a measure of impact (Fig. 4a; see Materials and Methods; Datasets S2 and S3). Using the so-called adstock function, we could account for the diminishing effect of each action in time. Our analysis revealed that direct contact was overall the most effective tool for recruitment across all regions of Europe (Fig. 4b). For every 100 people reached, an average of 1.2 participants registered (t = 5.730, p-value < 0.001). Each direct contact also required the presence of an outreach medium, and, in turn, outreach media required follow-up advertising. Yet, our analysis suggests that the impact of outreach media was crucial on its own, as it reached an average of 4.6 registered participants for every 1,000 people reached (t = 2.176, p-value = 0.0319). Outreach was particularly effective in Western Europe (Fig. 4b). Attending events such as forest fairs was also an effective way of recruiting: contacts made at events often led to registrations, particularly in Southern Europe (Fig. 4b; t = 2.084, p-value = 0.0397). Social media did not contribute to the recruitment (t=-0.958, p-value=0.3402) and even had an adverse effect in some regions (Fig. 4b). We hypothesize that this is because social media is based on the idea of regularly posting small amounts of information to keep followers, which is not suitable for advertising just once for an engaging project like MyGardenOfTrees.

**FIGURE 4.**
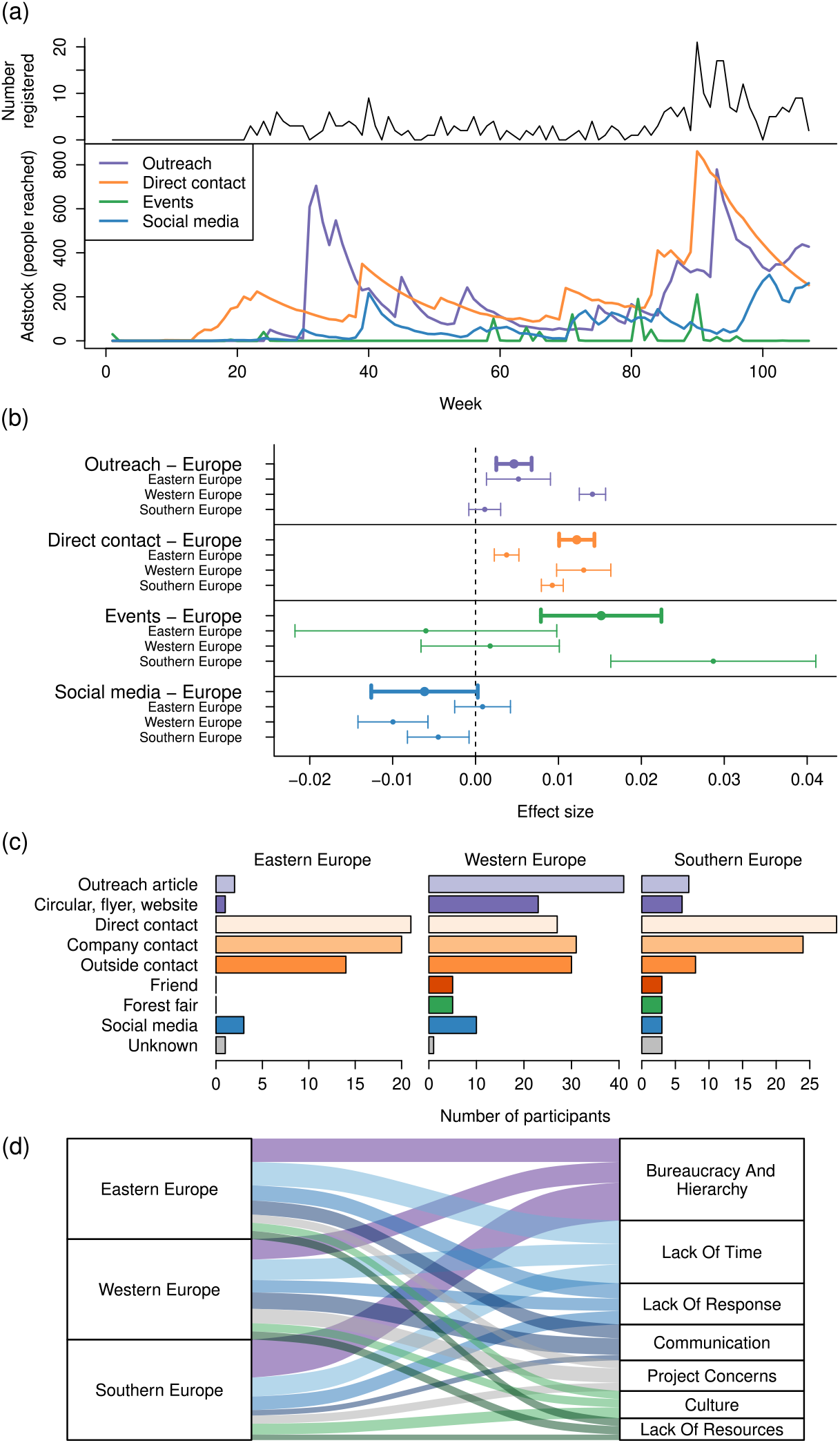
(a) Timeline of participant recruitment in MyGardenOfTrees across the 107 weeks: the top panel shows the number of registrations, while the bottom panel shows the number of people reached by the recruitment actions after applying the adstock function (see Materials and Methods). (b) Effect sizes were estimated for each recruitment action across Europe and within each region, using a marketing mix model approach. (c) Participant answers to the marketing attribution question of the registration form (Text S1). (d) Frequencies of coded interview sequences, with a focus on barriers to recruitment, indicated by the semi-structured interviews with the local coordinators.

The participants’ answers to the marketing attribution question (Question 17: “where you heard about MyGardenOfTrees?”, Text S1) provided independent evidence to our findings from the marketing analysis (Fig. 4c). It highlighted the role of outreach, particularly in Western Europe, and the crucial importance of direct contact. Specifically, we observed the impact of people who were not directly contacted by us spreading the word about the project (Fig. 4c): many joined not because of “direct contact” from a local coordinator, but because another person from the same or a different company, or even a friend, told them about the project.

Beyond understanding what worked, semi-structured interviews with local coordinators helped us identify the main barriers to recruitment across Europe (Fig. 4d; Materials and Methods; Text S3 and Dataset S4), as well as some regional differences (Kruskal-Wallis *chi*^2^ = 7.0777, df = 2, p-value = 0.029). The most frequently mentioned barrier, especially in Southern and Eastern Europe, was bureaucracy and hierarchy (Fig. 4d). Indeed, administrative and organizational obstacles within countries, as well as their public and private organizations, were common, including difficulties in reaching decision-makers, complex approval processes, and rigid hierarchical structures. The second most common obstacle was resistance from foresters to join due to a lack of time. Understandably, it is a challenge to commit to frequent monitoring over a five-year project duration. However, external circumstances, including the war in Ukraine, also contributed to the decision not to participate. On the local coordinators’ side, challenges in effectively conveying project fundamentals and identifying optimal recruiting strategies for each target group in a crosscultural context were reported (Fig. 4d). For example, one coordinator said, “I learned that every region and even province has its own way of forestry” (Dataset S4). Concerns related to the project itself, including its scientific objectives, were relatively rare (Fig. 4d), even though fears of introducing foreign genetic material and resistance from participants located outside the distribution range of the target species occurred. Indeed, the legislative landscape of Europe concerning the introduction of non-native tree species is complex and lacks transparency in some countries (Pötzelsberger et al. 2020). On the positive side, despite most local coordinators being female (Table S1) and forestry being a male-dominated sector in most countries, gender was never reported as a barrier.

### 3.3 Design and implementation of a continent-wide transplant experiment

MyGardenOfTrees is a distributed experiment designed to thoroughly explore the environmental gradients of interest through the appropriate randomization of experimental units (Fig. 5). This approach was ideally suited to overcome the difficulty that not all genetic units could be tested in all gardens, while ensuring all genetic units are represented across the same wide range of environmental gradients (Fig. 1a and b). The gray lines of Fig. 1c show that almost all seed families were represented across the full environmental gradient represented by the natural range of species and the seed origins. The representation of families was wider but less even for the precipitation gradient than in the temperature gradient due to higher variability across space, principally related to topography (Fig. 1c).

**FIGURE 5.**
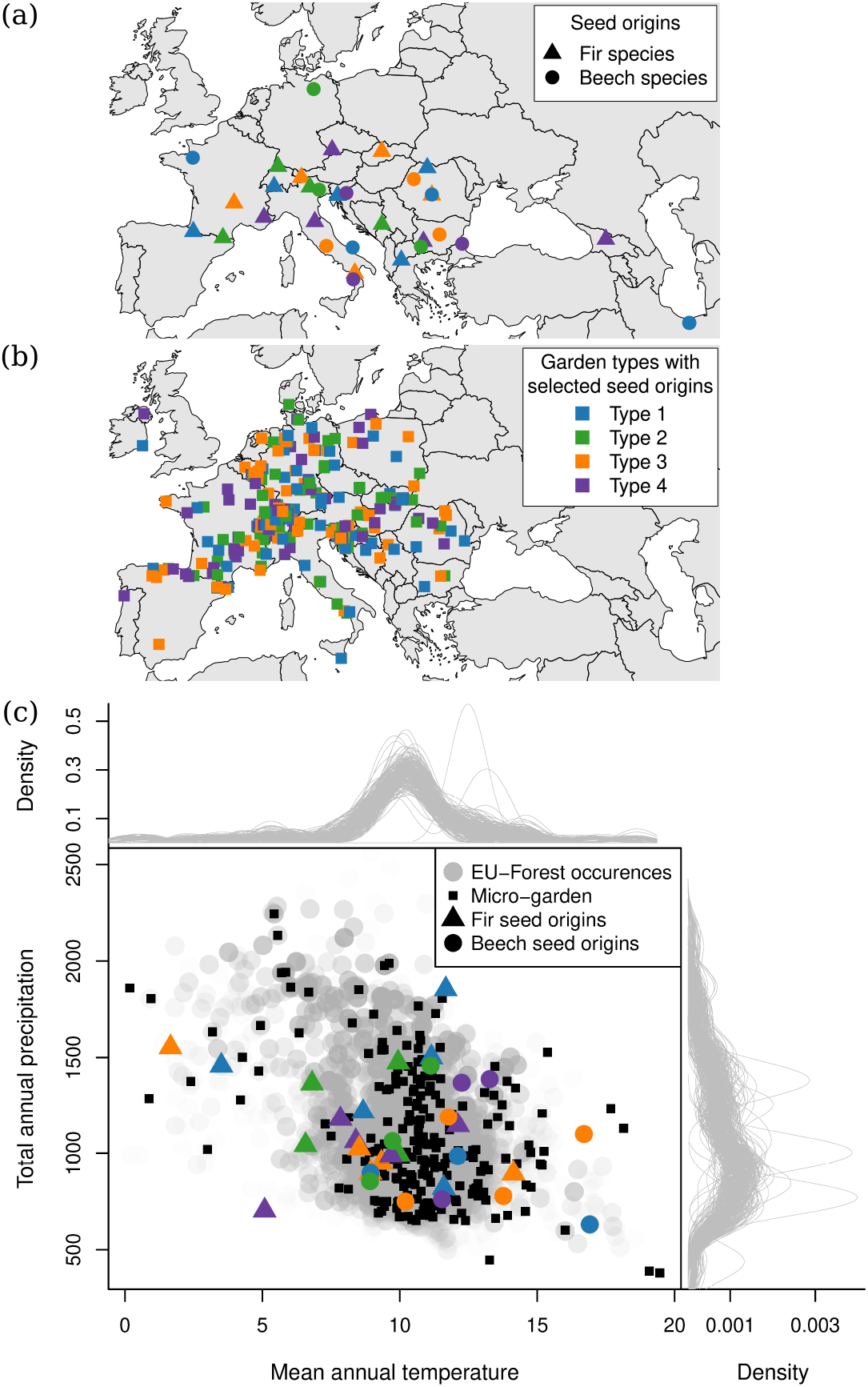
The realized experimental design of MyGardenOfTrees. (a) Seeds were sampled from 19 fir and 14 beech species and populations, with an average of 10 trees (families) per location. Only a selection of seed origins was distributed to each of the experimental gardens, defined by the type of garden. (b) The realized experimental design effectively represents the climate space of fir and beech forests in Europe (as defined by the EU-Forest database (Mauri et al. 2017)), and the tested 285 families (represented by gray density curves) are well-represented across the environmental gradients.

Foresters easily adhered to our complex protocols, which requested the selection of an appropriate forest area for the experiment, proper storage of the forest reproductive materials upon receipt, and sowing of seeds of 25 different origins with appropriate labeling (Text S2). Similarly, submitting observations about spring and autumn phenology, seedling survival, and damage was natural to them. Data collection, aggregation, and quality control were achieved using the open-source form Open Data Kit (ODK, https://docs.getodk.org) (Hartung et al. 2010) and smartphones. We used Ona (ona.io) as our server provider, which offers offline forms that can be completed using ODK Collect (Android) or the web-based Enketo tool (Android and iOS). An example data collection form for setting up the micro-gardens is provided (Dataset S5). The collection of data at a centralized server enabled the efficient tracking of participants’ work and correction of errors, as well as the creation of an R Shiny interface, where participants could track their observations in near real-time. The tools mentioned above have shown significant improvements even within the project timeframe, and we anticipate further developments.

## 4 DISCUSSION

### 4.1 Sound science with NGCS

In this paper, we advocate the use of NGCS for hypothesis-driven fundamental science as a tool for dealing with context dependency. While CS is increasingly employed in ecology for producing large, longitudinal datasets, these datasets are still seldom integrated in academic publications (Theobald et al. 2015, Johnston et al. 2023), and the use of CS in evolutionary biology is lagging. MyGardenOfTrees, along with advances in participatory breeding (Santamarina et al. 2025), challenges the common assumption that citizens cannot conduct experiments and engage in long-term monitoring to address fundamental ecological and evolutionary questions (Castagneyrol et al. 2023). We argue that CS is not merely a complement to hypothesis-driven approaches (Dickinson et al. 2010) and that NGCS can be a principal research tool for understanding mechanisms behind ecological patterns. In fact, several outstanding questions in ecology and evolution, coupled with ongoing societal challenges, could benefit from an NGCS approach. Beyond the study of local adaptation in various forest tree species that we have showcased, distributed experiments may also be used to test and compare reforestation methods, such as direct seeding or planting, as proposed in a CDE (Leverkus et al. 2021). Groom et al. (Groom et al. 2021) called for “next-level” citizen science to study species interactions with CS experiments, along with alien species management (van Rees et al. 2022, Di Febbraro et al. 2023). Research addressing the global concern arising around declining pollinator abundances could also be tackled using NGCS. Cultivating native plant species in urban gardens (Fukase and Simons 2016) and promoting high plant species diversity (Majewska and Altizer 2020) may foster pollinator activity. While CS initiatives are already being used to increase pollinator activity in private gardens and public green spaces, they remain local and correlative (Persson et al. 2023, Fonturbel et al. 2023). Finally, in the most advanced field of participatory breeding, Santamarina et al. (Santamarina et al. 2025) even proposes performing experimental evolution experiments to track variants driving co-adaptation in crop mixtures in collaboration with farmers.

A fundamental strength of NGCS is that, by involving professionals from the outset, it leverages the wealth of local knowledge and stakeholder perceptions, acknowledging them as key actors from the beginning. (Cornwall 2008). Working with participants who hold a tertiary level of education is recognized as an inherent inequality in benefit sharing in CS (Haklay 2018). NGCS overcomes this limitation by collaborating with professionals who can contribute meaningfully to the research content and enhance the integration of research findings into policy and action (Dunlap et al. 2021). Stakeholder involvement enhances the societal acceptance of findings, facilitates implementation, and improves trust and information flow between research and practice in the country (Sterling et al. 2017). For example, in Canada, it was found that AM was better accepted in regions where researchers invested in outreach (Peterson St-Laurent et al. 2018). In MyGardenOfTrees, we found that in some countries, notably in Slovenia, Croatia, and Hungary, more public owners joined the project than expected based on the proportion of publicly owned forests in these countries (countries above the line on Fig. 6b), suggesting trust in research and forest governance. Independent evidence, based on the proportion of certified Forest Stewardship Council (FSC) forest area in these countries (Maesano et al. 2018), also suggests a high level of trust in the public forestry sector in the countries mentioned above. A similar effect was expected in Baltic and Scandinavian countries and the UK; however, MyGardenOfTrees did not have participants in these countries. Finally, we also observed a bias towards participants who already own and/or manage a diverse forest (Fig. 6a), thus those who are already managing their forest with consideration for biodiversity. This finding raises the question of whether participating in CS projects would merely connect those who already know about the studied issues. Even so, is it our responsibility as researchers to arm these participants with more knowledge, thereby making their participation more likely to lead to transformative changes at the societal level.

**FIGURE 6.**
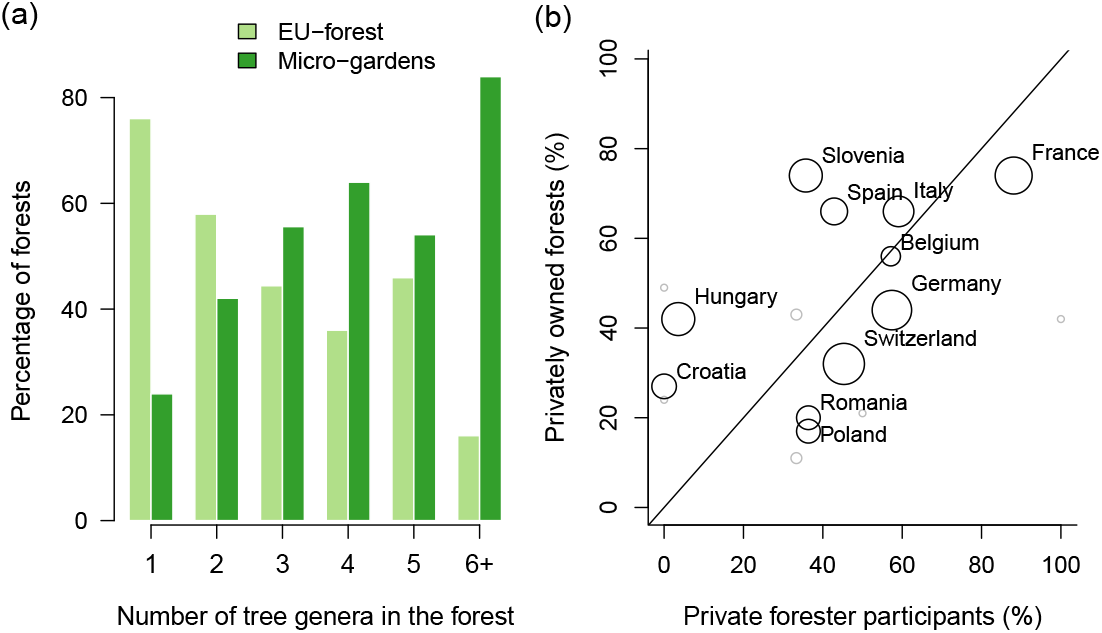
(a) The number of tree genera in the forested areas of Europe, according to the EU-Forest database, and the number of tree genera in the experimental micro-gardens of MyGardenOfTrees. (b) The percentages of privately (vs. publicly) owned forest areas across the European countries with at least five participants in MyGardenOfTrees, and the percentages of foresters with a private (vs. public) status who are participating in MyGardenOfTrees.

### 4.2 Recruiting and retaining participants

Dealing with context dependency by increasing the scale of experiments through NGCS has been demonstrated in agricultural and forestry contexts. However, challenges remain related to the human aspects and the characterization of the environment and ecologically relevant traits. First, large-scale and long-term experiments inherently involve linguistic and cultural barriers and challenges related to the allocation of resources and maintaining participant motivation. The allocation of resources to different recruitment strategies for future NGCS is a non-trivial question, and the academic literature lacks information about the factors that influence participation in CS projects (Nov et al. 2014, Bonney et al. 2014, West and Pateman 2016, Parrish et al. 2019). In marketing, saturation refers to the point at which increased media spending yields diminishing returns, meaning that each additional dollar spent yields smaller sales or engagement gains. In MyGardenOfTrees, direct contact with the target population, combined with outreach support items, appeared to be the best strategy. Thus, we recommend investing personnel and monetary resources in creating a few high-quality outreach support materials, such as a website, magazine article, or flyer, and supplementing them with direct contact via email or phone in the local language. It is also best if a locally respected person or authority delivers the message. Additionally, as others have recognized, respect for the volunteers and the country’s culture and administration is essential and should be studied before the recruitment process (Requier et al. 2020, West and Pateman 2016, Fraisl et al. 2022).

In CS or NGCS projects, as well as in CDEs, the work does not stop at recruitment but continues with efforts to motivate and retain participants (Parrish et al. 2019, Martellos et al. 2021), which raises challenges similar to those in CDEs (Fraser et al. 2013). Second, observations are limited to those visible with the naked eye, even though the installation of cameras or the analysis of posted samples (such as plant parts and feces) is also possible. Still, these aspects require the capacity of research labs to process a large number of samples. Connecting citizen observations with remote sensing is a promising avenue (Antonelli et al. 2023). Third, measuring relevant confounding factors locally and/or relevant aspects of the environment can be challenging because the number of observations must remain reasonable to avoid overburdening the participants. Interpolated climate, soil, and remote sensing data (satellite) are commonly integrated to replace local measurements. However, the importance of the microclimate is increasingly recognized (Kemppinen et al. 2024), calling for local measurements.

### 4.3 Gradient and distributed experiments with NGCS

Once participants are recruited, the implementation of a CS project is constrained by the capabilities, spatial location, and availability of participants. The most common study designs in ecological experiments, such as factorial, block, or split-plot designs, require a sufficient number of replicates for each group and treatment to facilitate the decomposition and comparison of variances within and between groups (Quinn and Keough 2002). Such designs are indeed difficult to achieve with CS, thus there is a major emphasis in the CS literature on these constraints (Johnston et al. 2023). It has only recently become apparent that some of these aspects, such as the dispersed location of participating citizens, can also facilitate hypothesis-driven experimental research. For example, participatory plant breeding leverages the expertise of farmers located throughout a country, and the heterogeneity in local farm conditions is ideal for breeding because farmers are also the end-users of the new varieties (Santamarina et al. 2025, Ceccarelli and Grando 2020, Brestovitsky and Ezer 2019, Danial et al. 2007). Similarly, in ecology, so-called gradient experiments that span a gradient of environmental conditions (instead of replicates) in the same environment (Tilman 1987, Kreyling et al. 2018) have been recognized as an efficient way to overcome context dependency (Catford et al. 2022). The use of many small experimental units that are placed across environmental gradients aligns exceptionally well with the concept of NGCS (Fig. 1a).

Today’s most popular type of CS project, targeting the general public at large spatial scales and without a specific protocol for data collection (Fig. 2c), can lead to data quality issues, survey inconsistencies, observer biases, communication issues, and other types of errors (Dickson et al. 2024, Bison et al. 2019, Dickinson et al. 2012, Balázs et al. 2021). Projects where participants receive in-person training (Tulloch and Szabo 2012, Fraisl et al. 2022, Johnston et al. 2023), where expert validation is incorporated, and where there is replication across volunteers, can partially resolve data quality issues (Danielsen et al. 2014, Golumbic et al. 2020). CDEs or NGCS projects leverage the potential of highly trained participants, potentially leading to more and higher data quality, thanks to the background knowledge of the participants. However, this approach will only be effective if there is an easy-to-use and transparent tool for data collection, aggregation, and quality control. In this context, the development of open-source form formats, such as the Open Data Kit (ODK, https://docs.getodk.org) (Hartung et al. 2010) has been key. ODK provides software and standards that underlie various solutions for form design and implementation, offering offline data collection, such as Enketo, Ona, and KoBoToolbox. ODK has revolutionized humanitarian data collection and enhanced clinical research in developing countries, but it also has emerging applications in agriculture (Ouma et al. 2019). Its applications in academic research are still scarce, where researchers often opt for standalone app development or CS dedicated platforms specific to image recognition/annotation, such as Zooniverse, or maps, such as SPOTTERON (see details in Liu et al. (2021)). The methods used in MyGardenOfTrees to develop the experimental design and observation protocols (Fig. 5, Text S2, Dataset S5) can provide guidance for future distributed experiments using both NGCS or CDEs.

## Abbreviations

CS: citizen science
CDE: coordinated distributed experiments
AM: assisted migration
NGCS: Next generation citizen science.

## AUTHOR CONTRIBUTIONS

K.C. obtained funding, designed the research, performed formal analysis, and wrote the first draft of the manuscript. M.B. performed formal analysis, contributed to the recruitment strategy of MyGardenOfTrees, and acted as local coordinator. N.P. contributed to the recruitment strategy of MyGardenOfTrees, coordinated the work of local coordinators, assembled recruitment data, and conducted semi-structured interviews. D.M.S. reviewed the CS literature, created the dataset of CS projects, and analyzed the semi-structured interviews. All co-authors contributed to the manuscript.

## ACKNOWLEDGMENTS

We are indebted to the 302 foresters who took part in this research, signed up for MyGardenOfTrees, and contributed numerous hours of fieldwork to this project. We also thank the multiple forest services and forestry companies of countries that made their participation possible. We thank all the local coordinators for their work, namely Sevil Cosgun, Marta Curman, Bernadett Fodor, Nancy Koller, Violeta Kotova, Mirjam Kurz, Azzurra Pistone, Bogdan Radu, Boris Rantaŝa, Camilla Stefanini, Fabian Suter, Karol Tomczak, and Maria Viota. KC thanks the ERC LS8 panel and the reviewers for believing that this project was possible. We thank Ona (ona.io) for their support of MyGardenOfTrees, and we acknowledge their excellent customer service.

## SUPPORTING INFORMATION

- Table S1: Profile of local coordinators responsible for recruitment and their countries of responsibility
- Text S1: MyGardenOfTrees registration form questions, including the informed consent form
- Text S2: General information sent to participants along with the “SetUpMyGarden” form
- Text S3: Semi-structured interview questions for local coordinators
- Dataset S1: Dataset of citizen science (CS) projects and coordinated distributed experiments (CDEs)
- Dataset S2: Recruitment actions of MyGardenOfTrees
- Dataset S3: Registrations to participate in MyGardenOfTrees (anonymised)
- Dataset S4: Coded segments of the semi-structured interviews with local coordinators – barriers
- Dataset S5: Code of the SetUpMyGarden Enekto XLSform

