## Supplementary Material for "Harnessing foresters’ engagement for climate change adaptation: the emerging tool of next-generation citizen science"

**This PDF file includes:**

Table S1: Profile of local coordinators responsible for recruitment and their countries of responsibility
Text S1: MyGardenOfTrees registration form questions, including the informed consent form
Text S2: General information sent to participants along with the "SetUpMyGarden" form
Text S3: Semi-structured interview questions for local coordinators

**Other supporting materials for this manuscript include the following:**

Dataset S1: Dataset of citizen science (CS) projects and coordinated distributed experiments (CDEs)
Dataset S2: Recruitment actions of MyGardenOfTrees
Dataset S3: Registrations to participate in MyGardenOfTrees (anonymised)
Dataset S4: Coded segments of the semi-structured interviews with local coordinators -- barriers

All SI datasets and analysis scripts are available at https://github.com/kcsillery/MyGardenOfTrees

**Table S1**: Profile of local coordinators responsible for recruitment and their countries of responsibility

| ID | Countries of responsibility | Languages spoken | Education | Occupation | Period of activity |
|---|---|---|---|---|---|
| 1 | Bulgaria | Bulgarian, Italian, English | Msc in environmental management | freelancer | September 2022 – 2024 |
| 2 | Croatia, Bosnia, Serbia | Croatian, Serbian, English | Msc in forestry | freelancer | May 2022 - Now |
| 3 | France, Switzerland (French speaking) | French, English | PhD in ecology | freelancer | April 2022 - Now |
| 4 | Germany, Austria | German, English | Forest engineer | freelancer | September 2022 - February 2024 |
| 5 | Hungary, Romania (Hungarian speaking), Bulgaria, Slovakia, Slovenia, Ukraine | Hungarian, English | Msc in Ecology | freelancer | April 2022 - Now |
| 6 | Italy | Italian, English | Msc in Ecology | PhD student in forest genetics | September 2023 - Now |
| 7 | Italy, Spain | Italian, Spanish, English | PhD in participatory science | project coordinator | April 2021 - March 2024 |
| 8 | Poland | Polish, English | Msc in forestry | PhD student in forestry | July 2022 - Now |
| 9 | Romania | Romanian, English | Bsc in forest sciences | Master student | May 2022 - |
| 10 | Slovenia | Slovenian, English | Msc in forestry | Associate at Forestry Institute and PhD student in Forestry | September 2023 - December 2024 |
| 11 | Spain | Spanish, English | Msc in environmental sciences | PhD student in environmental sciences | March 2023 – Now |
| 12 | Switzerland (German speaking), Germany | German, Swiss German, English | Bsc in forest sciences | Forest manager | October 2023 - Now |
| 13 | Switzerland (German speaking), Germany | German, Swiss German, English | Bsc in evolutionary genetics | Master student | June 2022 - December 2022 |

**Text S1**: MyGardenOfTrees registration form questions, including the informed consent form

### MyGardenOfTrees - Registration Form Trials 2023-2028

Welcome!

You are about to become a participant of MyGardenOfTrees. Before you fill in the following questions, make sure you have read the informed consent form.

#### Introduction

This questionnaire is aimed at clarifying information about your forest site where you would establish a micro-garden and obtaining your personal details in order to ship you the necessary materials. Before you fill in this questionnaire remember that:

1. By filling this questionnaire you will be officially considered as a participant of the MyGardenOfTrees project. This is why on the next page you will have to agree with an informed consent form.
2. By becoming a participant you commit to install and monitor a micro-garden for a minimum duration of 5 years (approximately 30 hours of voluntary work per year).
3. Retrieval from the project is possible at any time without consequences, nevertheless, we would like to minimize these cases because of the high cost associated to each micro-garden (>1000 EUR). Please make sure you have read the project expectations.
4. We assume that you have already identified a forest (subsequently called **Your forest**), where you have the permission to install the micro-garden either because you own and manage it or because you have obtained the relevant authorizations from the owner/manager. Note that you do not need to share this authorization with us but we may need to ask for it at any time.
5. MyGardenOfTrees micro-gardens are preferably installed in a forested area that is already fenced, can be fenced or where wildlife pressure is low.

If you need more information on what to expect, read here.
(https://www.mygardenoftrees.eu/future-participants/what-toexpect)

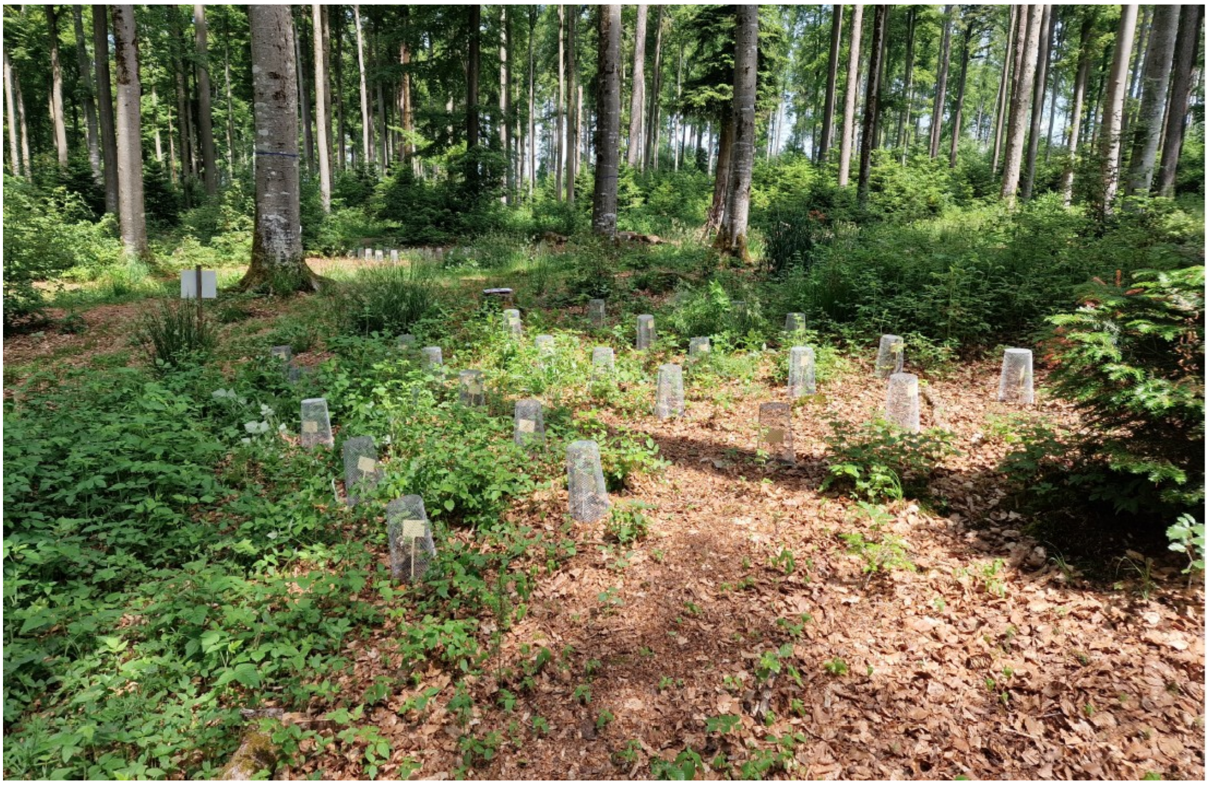

### Section 1: Informed Consent

**Question 1**: Please confirm that you agree with the Informed consent form by clicking on "Yes, I agree" below. If you do not agree or if you have questions, you can leave this form and contact us at

- [ ] Yes
- [ ] No

### Section 2: Forest Information

**Question 2**: What are the dominant and subdominant species of **Your forest**? (Check all that apply)

**CONIFERS:**

- [ ] Silver fir (Abies alba)
- [ ] Larch species (Larix spp.)
- [ ] Norway spruce (Picea abies)
- [ ] Swiss stone pine (Pinus cembra)
- [ ] Mountain pine (Pinus mugo/uncinata)
- [ ] European black pine (Pinus nigra)
- [ ] Scots pine (Pinus sylvestris)
- [ ] Douglas fir (Pseudotsuga menziesii)
- [ ] Common yew (Taxus baccata)

**BROADLEAVES:**

- [ ] Birch species (Betula spp)
- [ ] Hornbeam (Carpinus betulus)
- [ ] Black alder (Alnus glutinosa)
- [ ] Chestnut (Castanea sativa)
- [ ] European beech (Fagus sylvatica)
- [ ] Poplar (Populus spp.)
- [ ] Common oak (Quercus robur)
- [ ] Sessile oak (Quercus petraea)
- [ ] Turkey oak (Quercus cerris)
- [ ] Black locust (Robinia pseudoacacia)
- [ ] Lime species (Tilia spp)
- [ ] Elm species (Ulmus spp)
- [ ] Other: ________________

**Question 3**: What is the age of **Your Forest**?

- [ ] Less than 10 years
- [ ] Between 10 and 30 years
- [ ] Between 30 and 50 years
- [ ] Older than 50 years

**Question 4**: How is **Your Forest** managed?

- [ ] The forest has not been managed in the last 50 years and there is no plan of future intervention
- [ ] Continuous cover with an uneven-aged forest, natural regeneration only, cutting using single tree selection system
- [ ] Continuous cover with an uneven-aged forest, natural regeneration only, cutting using group selection system
- [ ] Even-aged forest with coppice or coppice-with-standards system
- [ ] Even-aged forest with shelterwood system
- [ ] A clearcut is planned but no sooner than in 6 years
- [ ] Other: ________________

**Question 5**: What is your relationship with **Your forest**?

- [ ] I own it and I manage it
- [ ] I own it, but I personally do not manage it
- [ ] I manage it and I have permission to use it for the 5 years of the trial from the owner (or the state)
- [ ] I have permission to use for the 5 years of the trial from the owner (or the state) and the manager
- [ ] I am planning to ask permission to use it

### Section 3: Micro-Garden Site Selection

**Question 6**: Have you identified the exact area in **Your forest** where you could establish Your future micro-garden?

*Remember that a micro-garden area is composed of 4 blocks of an area of 25 m² each. The 4 blocks can be dispatched in the forest but to facilitate the observations it is better to install them close to one another. Read here for more details about choosing the area: https://www.mygardenoftrees.eu/future-participants/what-to-expect*

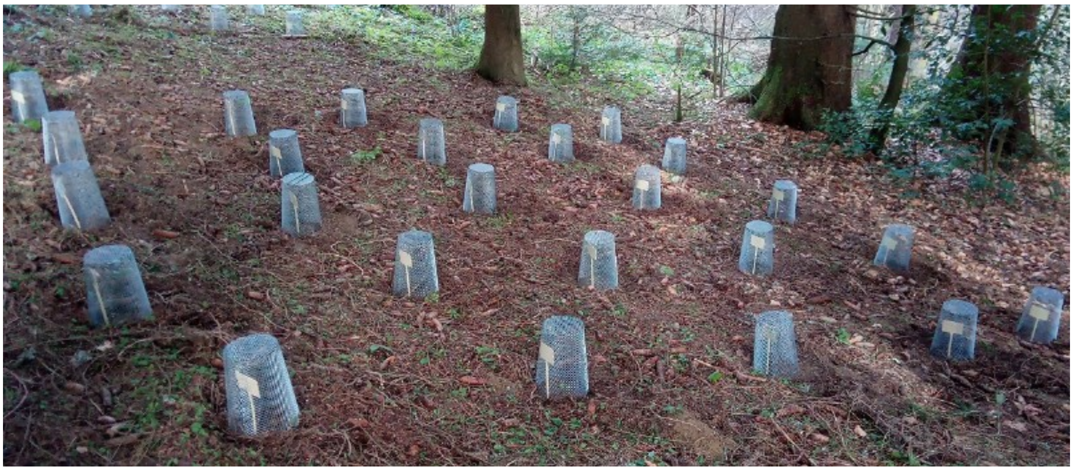

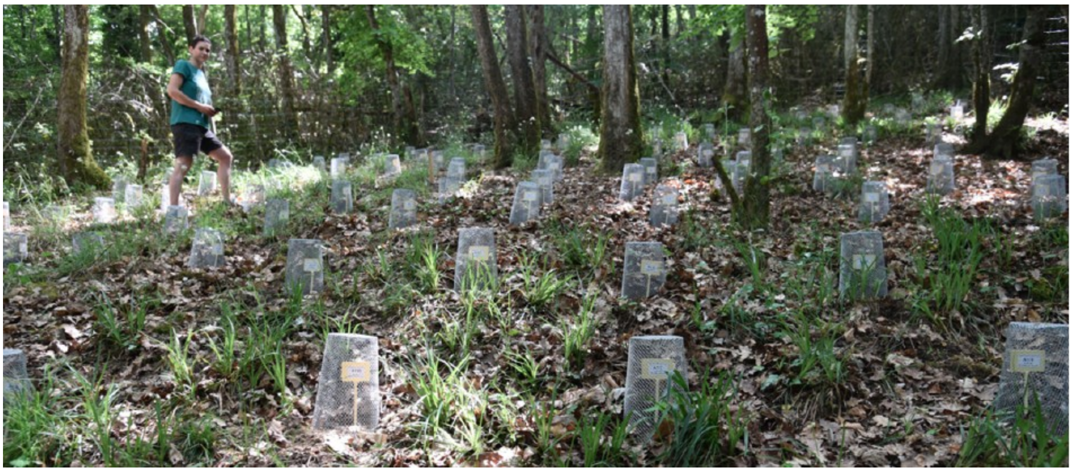

- [ ] Yes
- [ ] Not yet but the terrain is suitable for installing a micro-garden

**Question 7**: How quickly can you reach **Your forest** and **Your future micro-garden** from your home or workplace?

*Remember that each spring during 8 weeks, you will be asked to visit the site once a week to make observations that take up about 30 mins to 1.5 hours of your time, depending on the number of seedlings to be recorded, and excluding travel time.*

- [ ] In less than 30 mins
- [ ] Between 30 mins and 1 hour

- [ ] In more than 1 hour

**Question 8**: Is the area of **Your future micro-garden** fenced with deer proof fence?

*Note: Seeds and young seedlings will be protected by individual seed protectors. However, there is still a risk that wild boars could overturn the soil and the seed protectors, and/or that ungulates browse the young seedlings when the seed protectors will have to be removed after the 2nd or 3rd year of the experiment.*

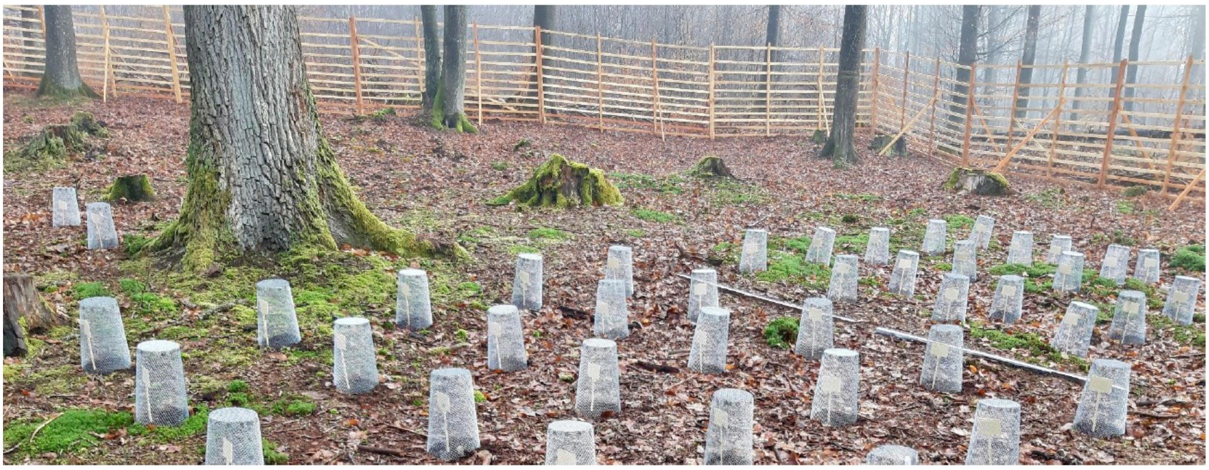

Please choose only one of the following:

- [ ] Yes
- [ ] No, but wildlife pressure is low
- [ ] No, but I could fence it at my own charge
- [ ] No, but I could fence it if I was reimbursed for the fencing materials
- [ ] No, and it cannot be fenced

### Section 4: Personal Information and Shipping

**Shipping Information**: We will ship you by post all the necessary material to install Your future micro-garden:

1. 100 seed protectors weighing around 30 kg
2. The seeds and the labels (seeds will need to be stored in a cool place below 10°C until sowing)
3. The protocols

Thereby we need to ask you some personal information. Remember that your personal information will only be used for shipping and will not be communicated to any third-parties (see Informed consent form that you agreed to).

**Question 9**: What kind of shipping address do you prefer to provide us? If you can, please provide us with both a private and a company or institutional address. This might be helpful for shipping reasons.

Please choose only one of the following:

- [ ] A private address
- [ ] A company/institutional address
- [ ] Both a private address and a company/institutional address

**Question 10**: What is your private shipping address? *All fields are mandatory. If you do not know how to answer some questions (e.g. additional information), please enter NA in that field.*

**Question 11**: What is your company or institution address? *All fields are mandatory. If you do not know how to answer some questions (e.g. additional information), please enter NA in that field.*

### Section 5: Participant Background

**Question 12**: Please tell us how would you like to be called? You can provide us either your full name, a nickname or a pseudonym that you like!

Please write your answer here:__________________

**Question 13**: Which of the following languages do you feel comfortable using? This will help us understand to how many languages we need to translate our protocols and other informative material. Please choose every language that you can comfortably use to read and communicate.

Check all that apply:

- [ ] Albanian
- [ ] Bulgarian
- [ ] Bosnian
- [ ] Croatian
- [ ] Czech
- [ ] Danish
- [ ] Dutch
- [ ] English
- [ ] Estonian
- [ ] Finnish
- [ ] French
- [ ] German
- [ ] Greek
- [ ] Hungarian
- [ ] Irish
- [ ] Italian
- [ ] Latvian
- [ ] Lithuanian
- [ ] Macedonian
- [ ] Norwegian
- [ ] Polish
- [ ] Portuguese
- [ ] Romanian

- [ ] Serbian
- [ ] Slovak
- [ ] Slovenian
- [ ] Spanish
- [ ] Swedish
- [ ] Ukrainian
- [ ] Other: ________________

**Question 14**: How many years of formal education have you received after the secondary school exit exam (for young adults between 15 and 20 years old)?

- [ ] 3 years or less
- [ ] More than 3 years

**Question 15**: Which was your main field of education after the secondary school exit exam?

**Question 16**: Is your primary occupation related to the forest sector?

- [ ] Yes
- [ ] No

**Question 17**: Finally, tell us where you heard about MyGardenOfTrees?

- [ ] Direct contact from the MyGardenOfTrees team
- [ ] From colleague inside the company/institution
- [ ] From a person outside my company/institution
- [ ] Social media ad posted by MyGardenOfTrees
- [ ] Circular/ad of another institution/association (e.g. EFI or IUFRO)
- [ ] Outreach article in the language of my country
- [ ] Other: ________________

### Section 6: Additional Comments

**Question 18**: Before you leave this questionnaire, do you have any question or comment for us?

---

**Thank you for becoming a participant of MyGardenOfTrees. We are looking forward to working with you!**

*The MyGardenOfTrees team*

---

**Text S2:** General information sent to participants along with the "SetUpMyGarden" form

MyGardenOfTrees - Trials 2023-2028

### How to set up your micro-garden?

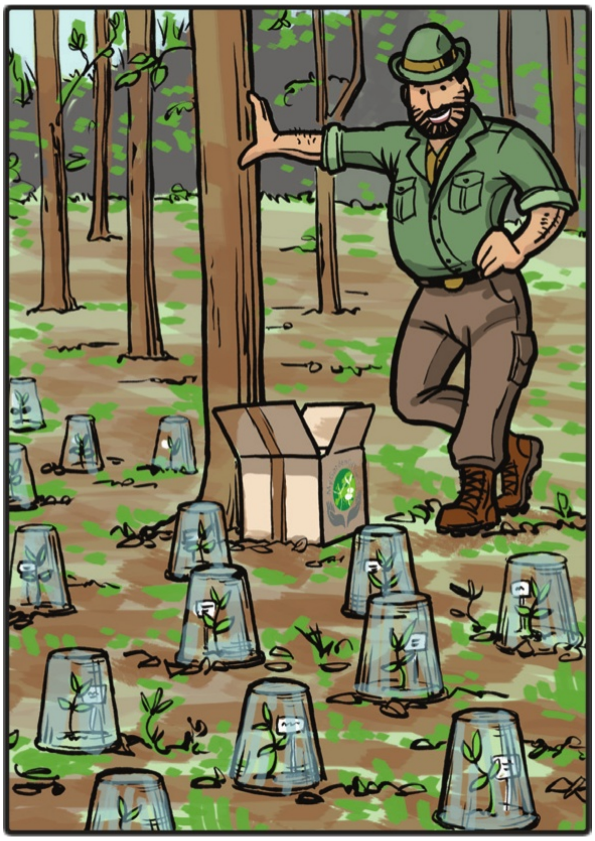

**©MyGardenOfTrees | Illustred by Sakál Design – *www.sakaldesign.com***

### 1. What to do with this parcel?

This parcel contains your seeds. The trials of MyGardenOfTrees mimic natural regeneration. This is why you received non-stratified seeds. We have stored these seeds at -5˚C since their collection in autumn 2022. Now it is important that you install the micro-garden as soon as you can so the seeds get a natural stratification typical to your forest. In the meanwhile, keep the parcel with the seeds in a cool and dry place. Ideally, you put the seeds in the plastic bags in your fridge (~5˚C) or in your garage or cellar. Make sure that you choose a place where temperature does not go above 10˚C. Nevertheless, if the maximum day temperature is below ~3-4˚C outside, we recommend that you wait for a warmer period to install your garden. The soil will be frosty and it will be difficult to handle the material and your smartphone. The seeds can last in the fridge up to some weeks.

### 2. Overview of your future micro-garden

##### 4 blocks spread out in your forest with at least 5m between them!

Your micro-garden is composed of 4 blocks**:** Block 1, Block 2, Block 3, and Block 4. The blocks can be spaced out in the forest but keep at least 5 m between blocks to make sure that you know which seeding spot belongs to which block!

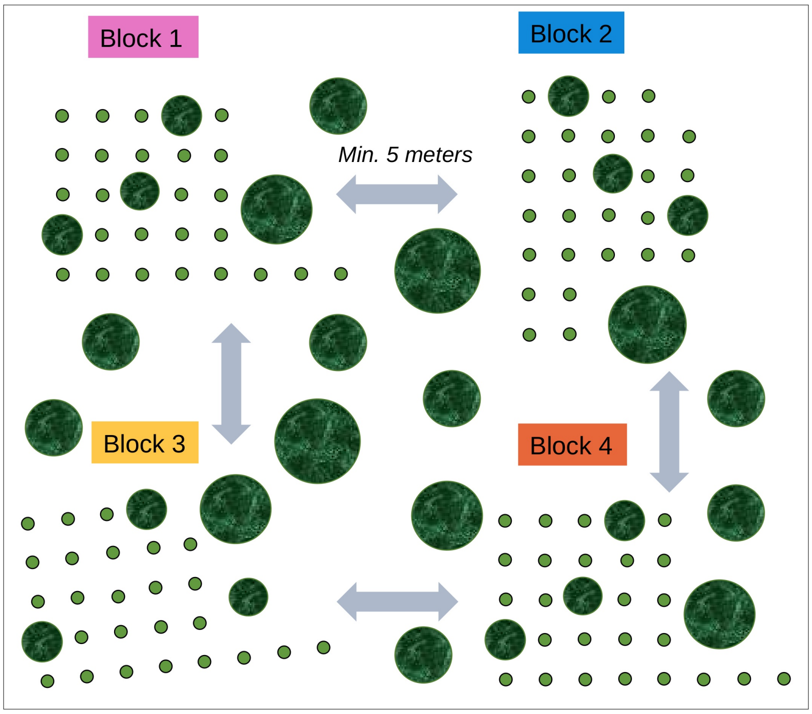


##### Zoom to a block: seed protectors must be in parallel lines!

Each block consists of 25 seeding spots. At each seeding spot, you will sow 20 seeds. You can arrange the seeding spots in any shapes depending on the obstacles you may encounter (such as trees, roots or stones) as long as you keep the seed protectors in parallel lines.

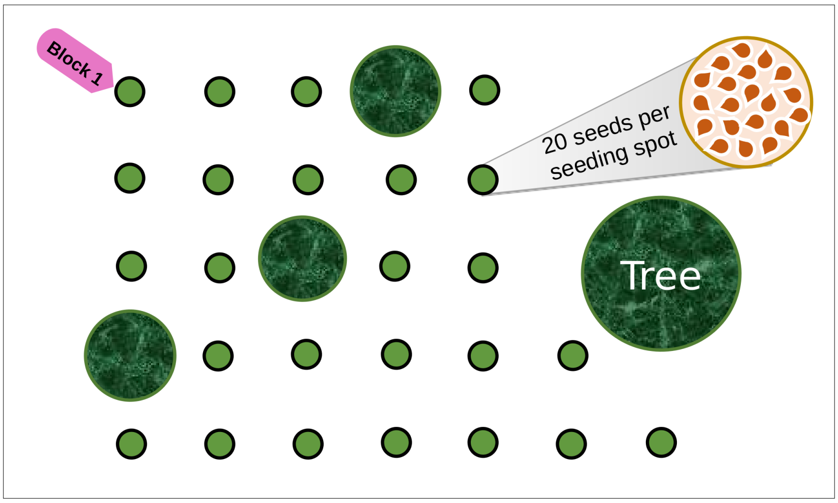

#### Each block has 25 different seed origins, but the 4 blocks are identical!

Within one block you will sow seeds from 25 different seed origins, i.e. species, provenances and families (mother trees), but the <u>4 blocks are identical, in fact, they are replicates</u>. You may sow the different seed origins in a different order across blocks.

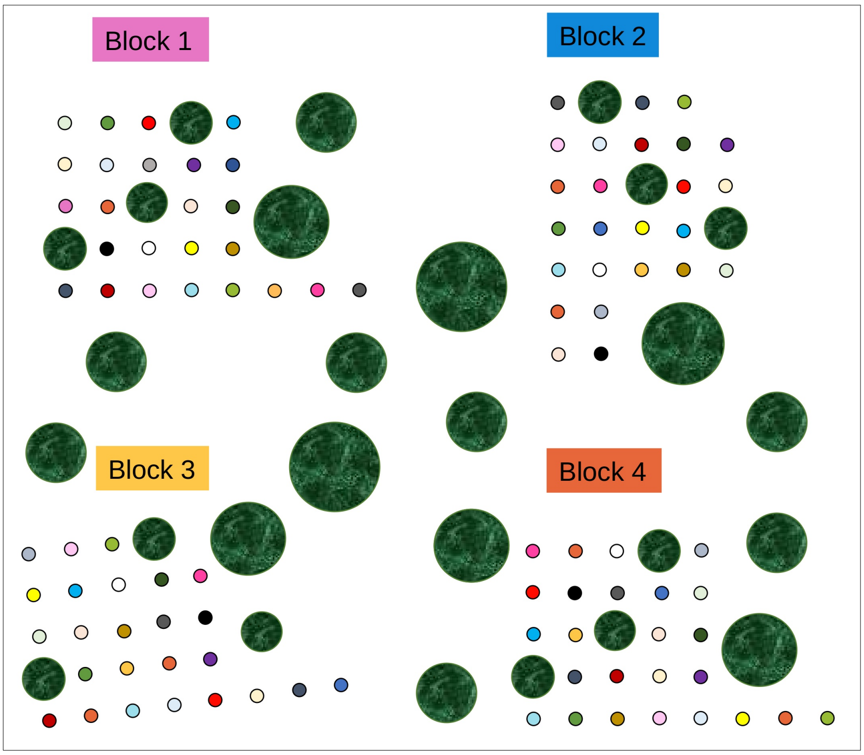


#### You will tell us the planting order of the seeds starting from the top left corner!

<u>You can choose a corner where you put the colored tag of the block</u> (pink for Block 1, blue for Block 2, yellow for Block 3, orange for Block 4). This will be the starting point to tell us the order in which you planted the different seed origins. Note that in the field, you use your smartphone and the <u>ENKETO Web Form</u> to tell the order!

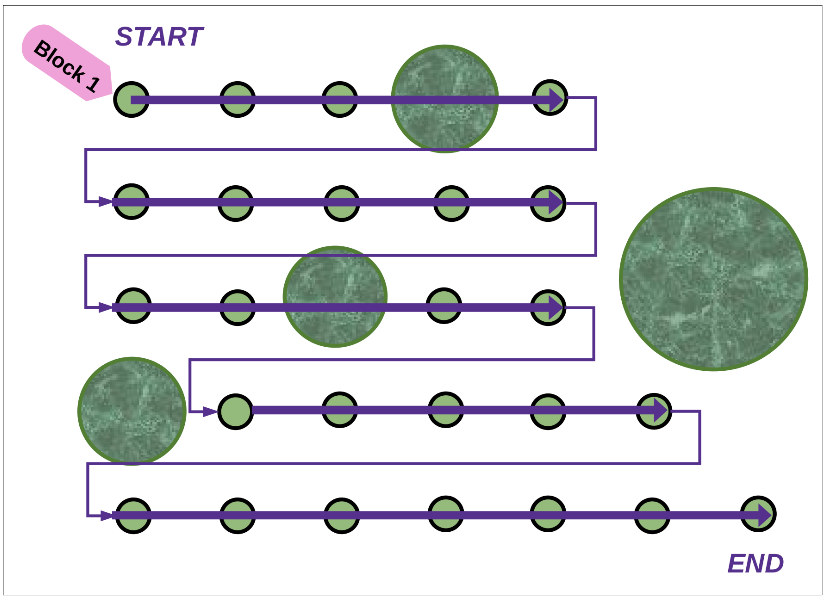


#### In the field, you will draw the shape of your block using the sheets with grids and send us a photo of it!

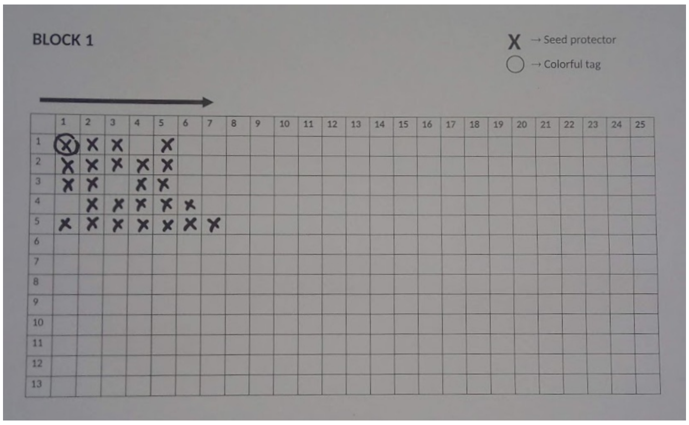

### The ENKETO Web Form

You will use your smartphone to collect and send us data about your micro-garden and the seedlings growing inside. We use a tool called “ENKETO Web Form”. To use it, you don’t need to install any apps!

To open an ENKETO Web Form for the first time, you need internet connection. Once opened, you can fill forms offline until you close your web browser[1]. Thus, if you have no or weak reception at your micro-garden: open the ENKETO Web Form with internet connection (at home) before going to your micro-garden!

#### Get to know the ENKETO web form “SetUpMyGarden”

| | |
|---|---|
| Insert this link https://enketo.ona.io/x/wocq0HYV in your mobile phone or scan the QR code. | |
| **Select your preferred language.** Click on the drop-down menu on the top to select the language you prefer.<br><br>While at home, **you can explore the form** to get comfortable with it. Don’t hesitate to fill in any information for testing. As long as you don’t submit the form, the information can be erased when you re-open the form. | 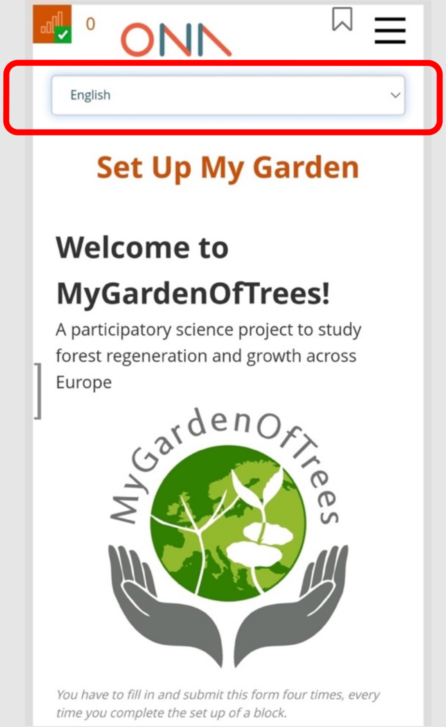<br> |

1 *Technically, until you clear your cache, but it is done automatically by Chrome and other web browsers (default option).*

<table>
<tr>
<td>By clicking on the brackets on the left of the screen, you can access the list of forms waiting to be submitted.</td>
<td>
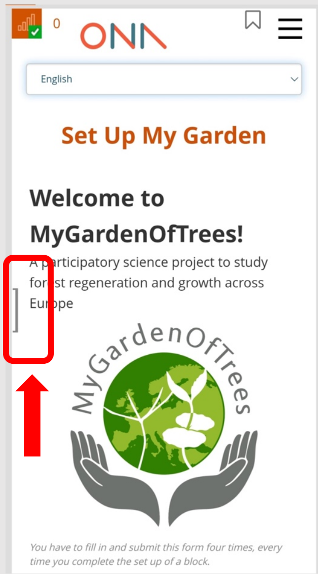

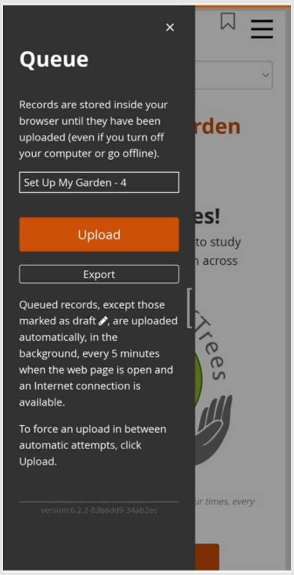

</td>
</tr>
<tr>
<td>On the top right corner, you can access the <u>full list of questions</u>. Click here in case you want to go back to a specific question to correct it or to explore the form.</td>
<td>
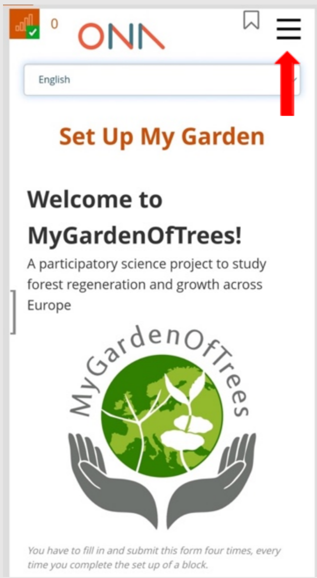

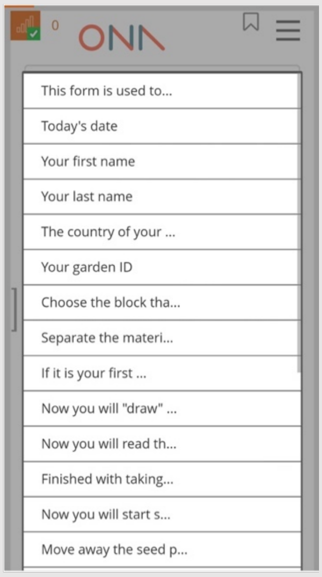

</td>
</tr>
</table>

### At home: How to get ready for installing your micro-garden?

| | |
|---|---|
| **Check the content of your parcels.**<br><br>Big parcels each contain:<br>• 50 metal seed protectors<br>• 150 metal pegs<br><br>Small parcel:<br>• 100 paper bags with seeds of *Abies* spp and *Fagus* spp. The seeds are delivered in 4 plastic bags, one for each block.<br>• 100 tags labeled with species name and provenance. The tags are stapled to the paper bags.<br>• 4 laminated warning signs, one for each block.<br>• A bag with 100 cable ties to attach the labels to the seed protectors.<br>• A white sheet of 1m by 1m.<br>• A 4m long string.<br>• A few empty white tags.<br>• 4 sheets with grids to draw the shape of each block | Two "**big parcels**": 50 x 50 x 50 cm parcels from LTM Italy<br><br>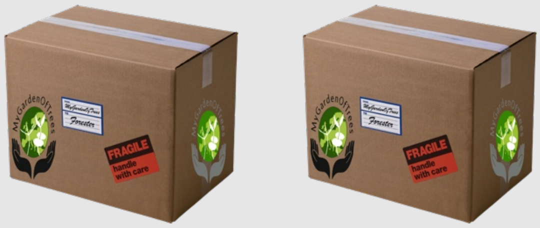<br><br>One "**small parcel**" 20 x 30 x 30 cm parcel from the WSL or your local coordinator<br><br>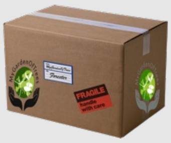 |
| **What else do you need to set up your micro-garden?**<br><br>• A pair of gardening gloves for handling the seed protectors.<br>• A hand fork or small rake to loosen the potentially frozen soil or to remove mosses and seedlings.<br>• A hammer to fix the seed protectors with pegs.<br>• A permanent marker to add your name and contact details to the warning sign if you wish to do so.<br>• 4 wooden poles and pins/nails to display the warning sign.<br>• A kneeling pad, or at least a thick plastic bag, to protect your knees! | 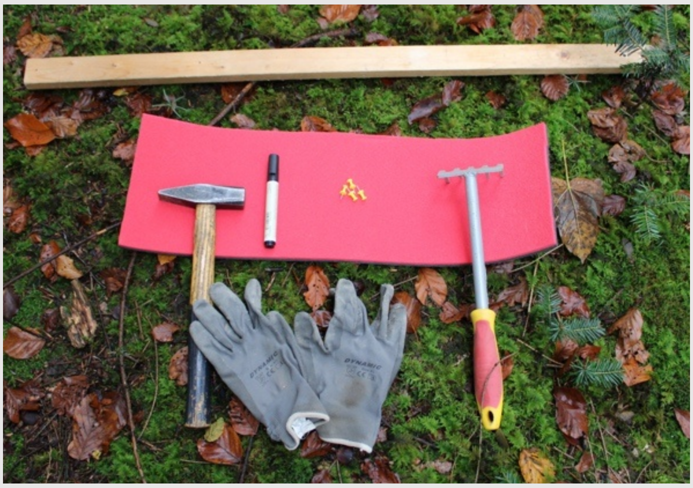 |

| | |
|---|---|
| **Fully charge your mobile phone before leaving to the field!**<br>• If you go with other people: ask everybody to fully charge their phones.<br>• If you have an external battery, take it for re-charge.<br>• Keep your phone in the pocket when not in use!<br>• If your phone runs out of battery and shuts down: don’t worry, your ENKETO Wed Form is not lost! When the phone is recharged, you can recover the form and continue to work with it. | 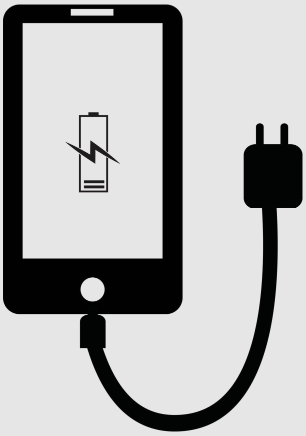 |

In order to enjoy your day of installing your micro-garden, we recommend that you wear warm and resistant clothes (clothes for forestry works are ideal)!

**Text S3:** Semi-structured interview questions for the local coordinators

**Overview**

Semi-structured interviews were conducted with all main local coordinators involved in participant recruitment for the MyGardenOfTrees citizen science project. This included coordinators responsible for single countries as well as those who worked across multiple national contexts.

**Interview Objectives**

The interviews were designed around three core objectives:

1. **Document recruitment strategies**: Capture the diverse approaches used by coordinators to engage potential participants across different countries and forestry sectors
2. **Identify barriers and facilitators**: Systematically catalog factors that hindered or facilitated successful recruitment in different contexts

**Interview Format**

- **Type**: Semi-structured interviews with open-ended questions
- **Duration**: Approximately 45-60 minutes per interview
- **Mode**: Conducted via video conference and audio-recorded with participant consent
- **Language**: English

**Interview Conduct**

- Interviews followed the structured question guide while allowing for natural conversation flow
- Follow-up questions were used to clarify responses and explore emerging themes
- Coordinators were encouraged to provide specific examples and case studies
- Sensitive topics (e.g., personal challenges, institutional conflicts) were handled with appropriate discretion

**Question Framework**

The interview guide consisted of 25 core questions organized into six thematic areas:

| **Question** | **Qualitative Theme** |
|---|---|
| 1. When did you join the project? | Which trials did they help with. |
| 2. How were you contacted? From whom? | Coordinators recruitment<br>Network base |
| 3. Why did you decide to join? | Network base<br>Level of commitment |
| 4. What did you think about the initial target of 500 participants? | Trust in the project and in the process;<br>Level of commitment |
| 5. What did you start with? Did we provide you with contacts? Did you have any communication material already available? | General communication strategy |
| 6. What was your initial recruiting strategy? Who did | General communication strategy |

| | |
|---|---|
| you target and through which channels? | |
| 7. What was the communication material did you use the most? | General communication strategy |
| 8. What were the first challenges and how did you adapt your strategy based on the challenge? | Barriers to recruitment |
| 9. What was the biggest challenge you encountered? | Barriers to recruitment |
| 10. What was the strategy or event which resulted in the highest number of participants? | Enabler |
| 11. Were you responsible for other countries? If yes, go through the same questions as above. | General communication strategy |
| 12. Think about a country where you tried to get participants, but you could not get any. What do you think were the main barriers? | Barrier to participation |
| 13. Did all participants from 2021-2023 re-registered for the new trials? If not, why? | Relationship between experiment success and participants engagement |
| 14. What did you miss which would have helped you significantly with the recruitment? | Possible future enablers |
| 15. What were the main reasons for participants not joining? | Barrier to participation |
| 16. What was the most common negative feedback from registered participants? | Barrier to participation |
| 17. What was the most common negative feedback from registered participants? | Possible barrier to monitoring/long term engagement to the project |
| 18. What was the most common positive feedback from registered participants? | Enabler |
| 19. Do you think your gender or ethnic origin had an impact on recruiting? | Enabler/barrier |
| 20. Do you think some of your character traits/personality affected the recruitment in some way? | Enabler/barrier |
| 21. What is your level of confidence with the scientific background of MGOT? Did it affect the recruiting process? | Enabler/barrier |
| 22. Can you think of an embarrassing moment you experienced during the recruiting process? | - |
| 23. What was the attitude toward the project in the public vs the private sectors? | Enabler/barrier |
| 24. Bottom-up vs top down recruitment. Which type did you use more, which one was more successful? | General communication strategy |
| 25. Overall, do you feel satisfied with your recruiting efforts? If you would start from scratch again, what would you change? | Possible future enablers |
